# Extracting and evaluating plant trait information from digital text in the Global Biodiversity Information Facility

**DOI:** 10.1101/2025.03.16.643436

**Authors:** Takuro Katori, Michio Oguro

**Affiliations:** Forestry and Forest Products Research Institute, Forest Research and Management Organization, 1 Matsunosato, Tsukuba, Ibaraki, 305-8687 Japan

**Keywords:** database, GBIF, growth form, life span, plant height, Raunkiæran shortfall, text mining, trait

## Abstract

**Premise of the study:** Plant traits are closely associated with species functions and environmental responses, and their compilation is particularly essential for large-scale studies. Although several databases of plant trait information have been published, comprehensive information on plant traits is lacking. To address this issue, additional easy-to-use data sources are required. This study examined digital text from descriptions of occurrences in the Global Biodiversity Information Facility (GBIF) as a novel source of plant trait information and evaluated its potential to mitigate existing gaps.

**Methods:** We focused on the digital text available from descriptions of occurrences in GBIF as a novel source of plant trait information. We collected information on life span, growth form, and maximum plant height for vascular plants from GBIF and other common trait databases. Using the resultant dataset, we compared the reliability (i.e., congruence of trait values in the focal database with that of the representative database) of trait values in GBIF with those from other databases and evaluated their novelty.

**Key results:** The trait information extracted from the GBIF exhibited reliability comparable to that of common plant trait databases. Additionally, the number of species with trait values increased 1.2–2.7 times when incorporating species data obtained solely from the GBIF with those from other databases.

**Conclusions:** Although digital texts in GBIF have not been previously used as a source of plant trait information, the results indicate that GBIF may be a valuable source of plant trait information.

## INTRODUCTION

Traits are morphological, physiological, or phenological characteristics that can be measured at the individual level (Violle et al., 2007). The diversity and distribution patterns of traits in individuals and species are driven by abiotic and biotic environmental factors. Consequently, trait information can be used to infer the ecological functions and environmental responses of species (Valladares et al., 2007; Ordoñez et al., 2009; Hoffmann and Sgrò, 2011; Foden et al., 2013), making the collection and publication of trait information crucial.

For vascular plants, a substantial amount of trait information has been collected and published (Wright et al., 2004; eFloras, 2008; Maitner et al., 2018; Kattge et al., 2020). Recent studies on plant traits have utilized data sources such as TRY (Kattge et al., 2011; Kattge et al., 2020), the Botanical Information and Ecology Network (BIEN; Maitner et al., 2018), eFloras (eFloras, 2008), the Rice database (the trait database constructed by Rice et al. (2019)), and published literature as sources of trait data. Several global-scale studies on plant traits have been published using existing trait databases and several other sources, revealing correlations between plant traits and the environment (Morin and Chuine, 2006; Lamanna et al., 2014; Šímová et al., 2018; Rice et al., 2019; Vallicrosa et al., 2022) and correlations or independence among traits (Wright et al., 2004; Morin and Chuine, 2006; Diaz et al., 2016; Pierce et al., 2017; Shiklomanov et al., 2020).

However, although these plant trait databases have been constructed, information regarding plant traits is limited (Maitner et al., 2022). This lack of plant trait information, also known as the Raunkiæran shortfall (Hortal et al., 2015), hinders the accuracy and comprehensiveness of research that requires substantial plant trait information (Cornwell et al., 2019; Perez et al., 2019; Maitner et al., 2022). Consequently, addressing this knowledge gap in plant trait information is a crucial challenge. Although trait information can be obtained from the literature or through direct measurements of preserved or live specimens, obtaining information for large numbers of species using such methods is challenging due to constraints of time, budget, and resources. To mitigate the lack of plant trait information, exploring digitized and publicly available sources of plant trait information that have not yet been utilized offers a promising alternative.

In this context, the GBIF (GBIF: The Global Biodiversity Information Facility, 2025) database is a potential resource that can be utilized to obtain plant trait information. GBIF has been primarily utilized in previous studies as a source of information on plant distribution (coordinates or locations) (GBIF Secretariat, 2021) and was classified as a database of “distribution” by König et al. (2019). However, GBIF also contains specimen images and text information regarding the individual and the surrounding environment, which can potentially serve as sources of trait information. Although the extraction of plant traits from specimen images on GBIF has been previously attempted (Kommineni et al., 2021; leaf length, width, and area), extracting information from texts has rarely been attempted. If the plant trait information available in the GBIF database is reliable and includes species that are not listed in common databases of plant traits (TRY, BIEN, eFloras, and Rice database), it could help mitigate the lack of plant trait information and enable comprehensive research.

In this study, we focused on vascular plants worldwide and investigated the novelty and reliability of plant trait information in the digital text available on GBIF to elucidate whether it could contribute to mitigating the lack of trait information. Specifically, we investigated 1) the reliability of the trait values of the same species between the GBIF text using a representative database as a reference. 2) the number of novel species for which trait information can be obtained by adding GBIF text as an information source compared to using only a common database for plant traits (TRY, BIEN, eFloras and Rice database)? Additionally, we removed species with suspicious trait values and evaluated their effects on novelty and reliability. Although GBIF text includes various plant traits, such as life span, growth form, plant height, leaf phenology, climbing habit, epiphytic habit, and flower color, in this study, we focused on life span, growth form, and plant height, which are particularly abundant in GBIF text.

## MATERIALS AND METHODS

### Overview of the analyses

We followed the data processing and analysis procedures described below and evaluated the reliability and novelty of the trait data obtained from digital text in the GBIF database. 1) In the initial step, we downloaded trait data from five databases and cleaned the data. 2) Next, as these databases can contain multiple records of a species, we constructed species-level datasets for each database. 3) To assess the reliability of trait values obtained from GBIF, we calculated the degree of congruency between these values and those from a reference database. We then compared the degrees of congruency of GBIF with those of the other databases. As the reference database, we selected eFloras, given that flora are typically compiled by experts and are expected to be highly representative (König et al., 2019). 4) To evaluate the novelty of the trait values obtained from GBIF, we counted the number of species with trait values not available in other databases. 5) Finally, we removed species with suspicious trait values from the datasets and examined their effects on the reliability and novelty of trait values for each database. All data processing and analyses were conducted using R software (version 4.1.1 (R-Core-Team, 2021), unless otherwise noted.

### Traits

We extracted and evaluated three plant traits: lifespan (*LS*), growth form (*GF*), and maximum plant height (*PH_max_*). These traits were selected on the basis of their functional and evolutionary significance. When grouping various plant traits based on their intercorrelations, they can be broadly classified into size-related and leaf economy-related traits (Diaz et al., 2016). *LS*, *GF*, and *PH_max_* are interrelated (Moles and Leishman, 2008; Diaz et al., 2016) and are classified as size-related traits, which are associated with various functional and evolutionary processes. For instance, size-related traits are linked to functions such as light competition, growth cost, water transport cost (Westoby et al., 2002; Moles et al., 2009), frost tolerance (Swenson and Weiser, 2010; Boonman et al., 2020), r-K selection (Diaz et al., 2016), proportion of polyploid species (Stebbins, 1971; Rice et al., 2019), evolutionary rates of molecular and ecological niches (Smith and Donoghue, 2008; Smith and Beaulieu, 2009; Hawkins et al., 2011), rates of speciation and extinction (Soltis et al., 2013), and range size (Morin and Chuine, 2006).

Although *LS* and *GF* classifications vary across databases, we reclassified trait values obtained from the databases into the annual and perennial categories for *LS*, and herbs and wood for *GF*. For this study, biennials were considered as annuals. Additionally, we assumed that the same taxon would not exhibit both annual and woody characteristics, similar to the findings of Rice et al. (2019). *PH_max_* values of each species and the values of plant height (*PH*) as *PH_max_* candidates were treated as continuous variables.

### The trait databases

The five databases used in this study exhibited varied content and structure, as summarized in Tables 1 and 2. The GBIF (https://www.gbif.org/) is a database of the occurrence of organisms worldwide. Although the GBIF is typically used as a source of distributional information, it also includes trait information in the descriptions of occurrences. The data can be downloaded in tabular format, with rows representing the occurrences of taxa (i.e., specimen or observation). We downloaded 337,549,876 vascular plant occurrences (GBIF.org, 2021, DOI: doi.org/10.15468/dl.qpuj6c, see the Data Availability Statement) for our study. Each occurrence (a single row in the downloaded table) was considered as one record. GBIF contains records written in several languages (Table 2). For the analysis, we e included only those in English, Spanish, Portuguese, French, German, Chinese, and Japanese.

**TABLE 1.** Summary of the contents and coverage of the databases used in this study.

| Database | Contents | Clade coverage | Geographic coverage |
| --- | --- | --- | --- |
| GBIF | • Taxonomy<br>• Distribution<br>• Traits (hardly used) | All organisms | World |
| TRY | • Traits | Plants | World |
| BIEN | • Taxonomy<br>• Phylogeny<br>• Traits<br>• Distribution<br>• Abundance | Plants | World |
| eFloras | • Taxonomy<br>• Traits<br>• Distribution | Plants | North America and China |
| Rice database | • Traits (only life span and growth form) | Plants | World |

**TABLE 2.**
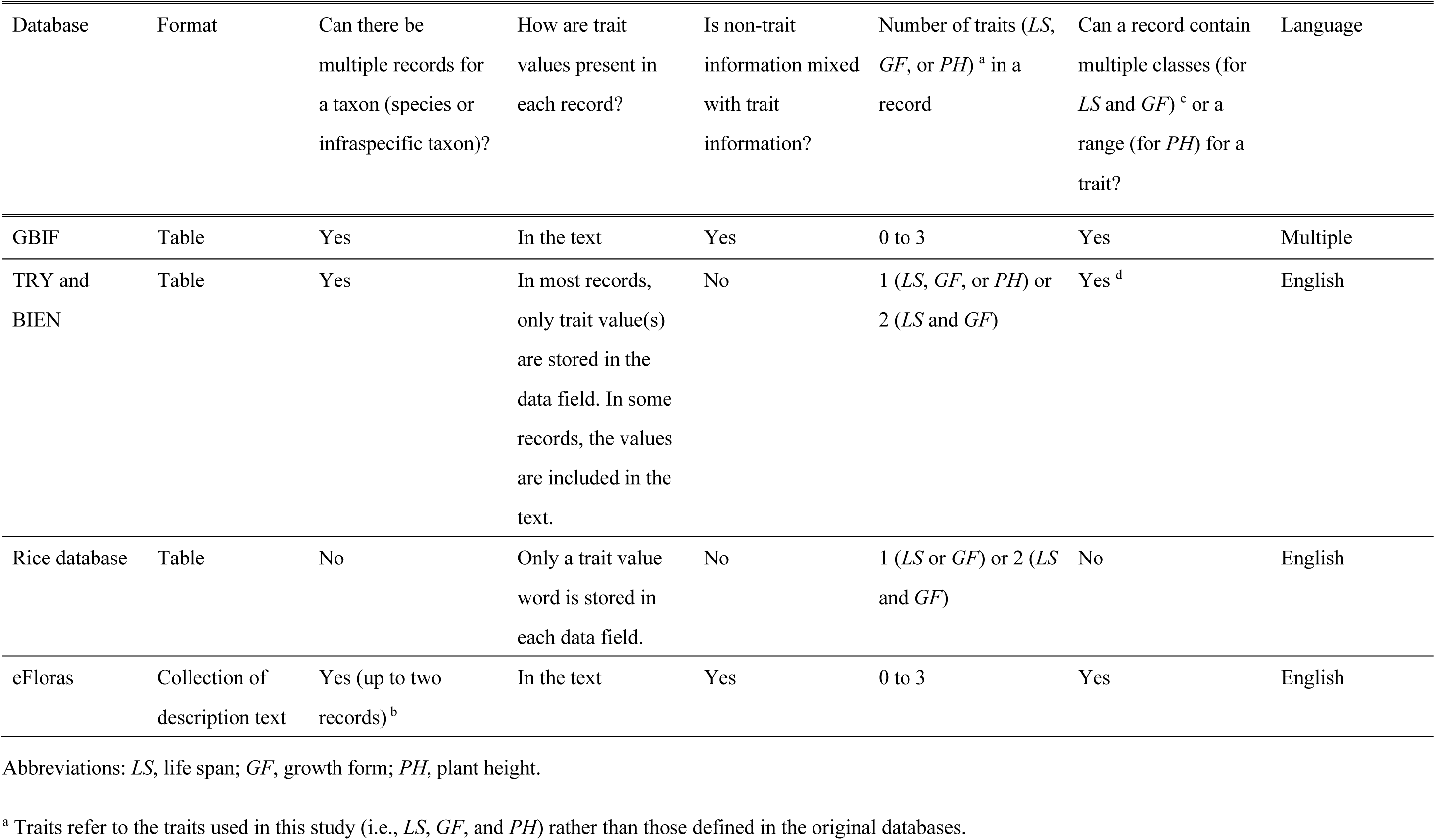
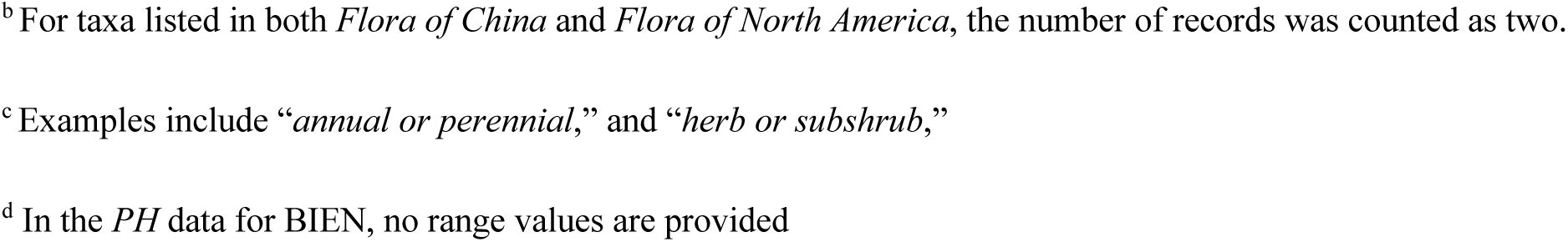
Overview of the structures of the trait databases used in this study.

| Database | Format | Can there be multiple records for a taxon (species or infraspecific taxon)? | How are trait values present in each record? | Is non-trait information mixed with trait information? | Number of traits (*LS*, *GF*, or *PH*)<sup>a</sup> in a record | Can a record contain multiple classes (for *LS* and *GF*)<sup>c</sup> or a range (for *PH*) for a trait? | Language |
| --- | --- | --- | --- | --- | --- | --- | --- |
| GBIF | Table | Yes | In the text | Yes | 0 to 3 | Yes | Multiple |
| TRY and BIEN | Table | Yes | In most records, only trait value(s) are stored in the data field. In some records, the values are included in the text. | No | 1 (*LS*, *GF*, or *PH*) or 2 (*LS* and *GF*) | Yes<sup>d</sup> | English |
| Rice database | Table | No | Only a trait value word is stored in each data field. | No | 1 (*LS* or *GF*) or 2 (*LS* and *GF*) | No | English |
| eFloras | Collection of description text | Yes (up to two records)<sup>b</sup> | In the text | Yes | 0 to 3 | Yes | English |

Abbreviations: *LS*, life span; *GF*, growth form; *PH*, plant height.
<sup>a</sup> Traits refer to the traits used in this study (i.e., *LS*, *GF*, and *PH*) rather than those defined in the original databases.
<sup>b</sup> For taxa listed in both *Flora of China* and *Flora of North America*, the number of records was counted as two.
<sup>c</sup> Examples include “*annual or perennial*,” and “*herb or subshrub*,”
<sup>d</sup> In the *PH* data for BIEN, no range values are provided

TRY (Kattge et al., 2011) is the most widely used database of plant traits worldwide. Between 2007 to 2019, it has been cited in over 250 publications (Kattge et al., 2020) and referenced in recent large-scale studies on plant traits (Sandel et al., 2015; Diaz et al., 2016; Butler et al., 2017; Pierce et al., 2017; Bruelheide et al., 2018; Cornwell et al., 2019; Perez et al., 2019; Boonman et al., 2020; Shiklomanov et al., 2020; König et al., 2021). TRY data can be downloaded in tabular format, with rows representing the value(s) of a trait for a taxon. We downloaded a table with 28,659,648 rows by selecting the relevant trait names from the website (https://www.try-db.org/TryWeb/Home.php; downloaded on November 27, 2022). The traits used for the analysis included *plant lifespan (longevity), plant growth form, Plant life form (Raunkiaer life form), plant woodiness, plant vegetative regeneration capacity*, and *plant functional type (PFT)* for *LS* and *GF*, and *plant height vegetative* and *plant height generative* for *PH*.

BIEN is a recently developed database that includes information on the taxonomy, phylogeny, traits, distribution, and abundance of plants worldwide (Maitner et al., 2018). It has been used for large-scale studies on plant traits (Kelly et al., 2021; Kling and Ackerly, 2021), and has a data structure similar to that of TRY. We downloaded a table with 10,619,951 rows by specifying trait names using the *BIEN_trait_trait* function in the R package *BIEN* version 1.2.4 on November 16, 2022. The trait names specified for the function were *longest whole plant longevity*, *maximum whole plant longevity*, *whole plant growth form*, *whole plant growth form diversity*, and *whole plant woodiness* for *LS* and *GF*, and *maximum whole plant height* and *whole plant height* for *PH*.

The Rice database (*Lifeform database* by Rice et al. (2019)) is a table listing *LS* and *GF* for plants worldwide. It was constructed by combining information obtained from multiple sources (eFloras, Encyclopedia of Life, Index to Chromosome Number in Asteraceae, International Legume Database & Information Service, Missouri Botanical Garden, World Checklist of Selected Plant Families, and a database compiled by Zanne et al. (2014)) and has been used for large-scale studies on plant traits (Poppenwimer et al., 2023; Wang et al., 2023). This database was obtained from the online supplementary information provided by Rice et al. (2019).

eFloras (http://www.efloras.org/) is an online database of flora from several countries and regions that contains descriptions of taxonomy, traits, and distributions. It has also been used in large-scale studies on plant traits (Thomson et al., 2011; Chen et al., 2019). Given that a machine-readable HTML format, rather than a PDF format, is available, we used *Flora of North America* and *Flora of China* in this study. We downloaded the description of each taxon from the two resources through web scraping using the R package *rvest* version 1.0.2, on August 26, 2023. Although the Rice database contains trait information obtained from eFloras, we also used eFloras as a source of trait information, as we believed that a higher number of trait values could be obtained from eFloras. This approach was selected because the Rice database only selected trait values directly described in the descriptions of species or infraspecific taxa, whereas Coleman et al. (2023) demonstrated that shared traits of a taxonomic group are commonly described in the description of the taxonomic group rather than the description(s) of each sub-taxon, and supplementing trait values from higher taxonomic groups can improve the availability of trait data. For this purpose, we obtained data not only from species and infraspecific taxa, but also from higher taxonomic groups such as families, genera, and sections.

### Extracting trait values from each record

Because databases other than the Rice database can contain multiple candidate values of *LS* and *GF* within a record (Table 2), we extracted all candidate values for each trait for each record from all databases. However, all databases include only one value of *PH* for each record. The sources of information for each trait in each database are listed in Table S1 in Appendix S1.

Because trait information in GBIF and eFloras is embedded in the descriptive text of occurrences/taxonomic groups, we conducted text processing, which searches for expressions representing trait values and assigns candidate trait values for each record. GBIF contains records written in several languages (Table 2). Among them, we prepared expressions intended to target trait values written in English, Spanish, Portuguese, French, German, Chinese, and Japanese; however, they could occasionally match words written in other languages, including those other than trait values. Although TRY and BIEN are structured databases, the values of *LS* and *GF* are not expressed as predefined classes; in some cases, these values are expressed as text. Therefore, we extracted the candidate values of *LS* and *GF* in TRY and BIEN by applying a text-processing procedure similar to that of GBIF and eFloras. Before extracting the trait values, all letters and numbers were converted into half-width lowercase letters without accent marks. Additionally, plurals were converted into singulars for languages other than German, Chinese, and Japanese.

#### Life span and growth form

To extract candidate trait values from the records in GBIF and eFloras, we searched for expressions representing each class of *LS* and *GF* from the texts pertaining to occurrences and taxa. However, these texts commonly contain expressions that can result in false detections of trait values (e.g., *annual rainfall*). Therefore, we removed these expressions from all records obtained from GBIF and eFloras before trait value extraction. For this purpose, we created a list of such expressions by searching for words and phrases representing trait values in randomly selected records and recording expressions that potentially disrupt their extraction. A list of these expressions is provided in Table S2. After excluding these expressions, we searched for expressions representing values of *LS* and *GF* shown in Tables S3 (GBIF) and S4 (eFloras) and obtained candidate values for each trait.

As noted in “The trait databases” section, the shared traits of a taxonomic group are typically listed in the description of the taxonomic group rather than in the description(s) of each subtaxon in the flora (Coleman et al., 2023). Therefore, for eFloras, we extracted candidate values of *LS* and *GF* from records of higher taxa, including family, genus, section, and species, of the same flora and assigned them when the values (s) of *LS* or *GF* were missing for terminal taxa, such as species or infraspecific taxa.

Candidate trait values in TRY were extracted using combinations of values in the three columns (*TraitName*, *DataName*, and *OrigValueStr*). Referencing these columns, we assigned candidate trait values to each record. The combinations of values and the corresponding trait values are listed in Table S5. Similarly, candidate trait values from BIEN were extracted and assigned to each record using a combination of the values in the two columns (*trait_name* and *trait_value*; Table S6).

As the trait values in the Rice database are classified in a manner similar to that in this study, they were not altered. Unresolved values (*mixed*, *conflict*, and *NA*) were regarded as missing data.

After obtaining the candidate values of *LS* and *GF* for each record in each database, we determined the final values of *LS* and *GF* for each record. Because we assumed that all annual species were herbaceous and all woody species were perennial, conflicts were resolved and missing information was complemented for each record using the following steps: 1) When a record contained annual and wood values, *LS* and *GF* were regarded as missing values, and steps 2–5 were omitted. 2) When a record contained values for herb and wood, *GF* was regarded as a missing value and Step 3 was omitted. 3) When a record contained values for wood, a perennial was used as the final value of *LS* for the record. 4) When a record contained annual and perennial values, a longer *LS* (i.e., perennial) was used as the final value and step 5 was omitted. 5) When a record had annual value and lacked perennial value, *GF* was regarded as an herb. For the other cases, the candidate values of a record were used as all conflicting classes had to be resolved using the steps above.

#### Plant height

For GBIF and eFloras, we obtained values of *PH* from the description texts of occurrences/taxonomic groups by matching patterns of expressions representing information regarding *PH*. To simplify the patterns, we initially removed expressions which could obscure the patterns prior to extracting *PH* values from these databases. These expressions included mathematical signs (>, <, and ±) and words, and signs represent approximation which are placed immediately before numbers (e.g., *ca*; all expressions are listed in Table S7). Additionally, we created a list of expressions that could result in false detection of *PH* values using the same method employed for *LS* and *GF* (Table S8), and all expressions on the list were removed prior to extraction. Finally, we matched the patterns of expression (Text S1) representing information regarding *PH* and extracted the value of *PH* for each record. For TRY, the values and units for *PH* were obtained from the *StdValue*, *OrigValueSt*, and *UnitName* columns, placing a higher priority on values in the *StdValue* column. It is important to note that *PH* values in TRY are occasionally expressed as ranges. For BIEN, values of *PH* were obtained from the *trait_value* column.

When the value of *PH* was recorded as a range (applicable for GBIF, eFloras, and TRY; symbols used to represent ranges included −, *∼*, *a*, and *to*; e.g. *1-2m*, and *1m-2m*, among others), the upper limit of the range was used as the *PH* value of the record. All *PH* values were converted to meters.

### Standardization of taxon names

To obtain a species-level dataset, we extracted taxon names (species or infraspecific taxa) from each record in each database. The locations used to extract the taxon names were as follows: GBIF, *acceptedScientificName* column; TRY, *AccSpeciesName* column; BIEN, *scrubbed_species_binomial* column; the Rice database: *Species* column; eFloras: top of each description text. The synonyms included in the taxon names were standardized. For reference, we used the Leipzig Catalog of Vascular Plants (LCVP, Freiberg et al., 2020). Standardization was performed using the *lcvp_search* function in the R package *lcvplants* version 2.0.0. Following standardization, infraspecific taxon names were converted to their type species. When a taxon was standardized to a taxon higher than a species (e.g., genus) or standardization was unsuccessful, the record was excluded from the dataset.

### Exclusion of PH value outliers through known maximum records

To reduce the potential effects of outliers on *PH*, we removed *PH* values higher than the known maximum records for higher clades (monocots, ferns, and other plants) in the existing literature. The maximum records in the literature are: 20 m for ferns (Wee et al., 2014); 61 m for monocots (Bernal et al., 2018); 116.07 m for *Sequoia sempervirens* (Sillett et al., 2021); and 100.8 m for other plants (Shenkin et al., 2019; Sillett et al., 2021). Using these maxima, *PH* values higher than the known maxima for each clade were removed. The higher clade name of each species other than *Sequoia sempervirens* was obtained by attaching order names through the taxonomic standardization described above, and by attaching class names using the species matching tool provided by GBIF (www.gbif.org/tools/species-lookup). We classified Liliopsida as monocots, Polypodiopsida and Lycopodiopsida as ferns, and other classes as other plants.

### Construction of species-level datasets for each trait for each database

Given that databases can contain multiple records for a species (Table 2) and different species may be merged into a single species through taxon name standardization, a trait of a species in a database may have multiple records with different trait values. Therefore, the representative value of each species for each trait in each database was determined as follows, and the resulting dataset was uploaded to Zenodo (see the Data Availability Statement):

#### GBIF, TRY, and BIEN

For *LS* and *GF*, we used the more frequent class for a trait (annual or perennial for *LS*; herb or wood for *GF*) as a representative value for each species in each database. When both classes contained the same number of records, the trait values of the species in the database were considered missing. For *PH_max_*, the representative value for each species in each database was determined as follows: Initially, to remove outliers in *PH*, the Smirnov-Grubbs test (Grubbs, 1969) was repeatedly applied for each species until no outliers were found. Next, using the maximum value of the remaining *PH* values of the species, we removed outlier species for each genus using the method described above. The maximum value of *PH* for each remaining species was considered a representative value.

#### The Rice database and eFloras

The Rice database and eFloras contain a limited number of records for each species; the original number of records for each taxon was limited, with only one record obtained for the Rice database and one or two records obtained for eFloras (Table 2). Therefore, we excluded species with both classes of candidate values for *LS* or *GF* from the dataset for that particular trait. Similarly, we determined a representative value for *PH_max_* by calculating the maximum *PH* value of the records for each species in each database without excluding outliers.

### Reliability of plant traits in GBIF

To determine whether the trait values extracted from GBIF had comparable quality with those of the other databases, we compared the reliability of the trait values obtained from GBIF with those of other databases by checking the congruency of the trait values with the reference database, eFloras. To achieve this, we defined the reliability indices as follows: For *LS* and *GF*, the reliability index was defined as the ratio of species with trait values congruent with eFloras to the total number of species with a focal trait value in the focal database. We assumed that a higher index indicated higher reliability of the trait in the database. For *PH_max_*, the reliability index was defined as the mean absolute value of the log_10_ *height ratio*, where the *height ratio* was determined by dividing the *PH_max_* of a species in the focal database by the *PH_max_* of the same species in eFloras. We assumed that a lower mean absolute value of the log10 *height ratio* represented a higher reliability of *PH_max_* in a database. Using these definitions, we calculated the reliability indices for each trait of GBIF, TRY, and BIEN. The Rice database was excluded from the evaluation as it contains data quoted from eFloras.

### Novelty of plant traits in GBIF

The novelty of the trait values obtained from GBIF was evaluated by counting the number of species with trait values that could be obtained from GBIF but were missing from the other four databases (TRY, BIEN, Rice database, and eFloras). To achieve this, all described species of vascular plants (351,180 species, Freiberg et al., 2020) were divided into the following four groups, and the number of species in each group was counted: a) species with focal trait values available only in databases other than GBIF, b) species with focal trait values in both GBIF and the other four databases, c) species with focal trait values only in GBIF, and d) species for which focal trait values could not be obtained using any of the databases (a + b + c + d = 351,180). Each trait was analyzed separately. We then evaluated the novelty of GBIF (the extent to which GBIF could mitigate the lack of trait information) by comparing the proportion of the number of species with trait values in databases other than GBIF to the total number of species ((a + b) / 351,180), and the proportion of the number of species with trait values in all databases, including GBIF, to the total number of species ((a + b + c) / 351,180) for each trait.

### Removal of species with suspicious representative value

We tested whether the removal of species with potentially incorrect trait values could improve the reliability of the trait values for each database. To achieve this, we defined the criteria for species removal and removed species by applying threshold values. For the removal of *LS* and *GF* values, we used the proportion of the representative values, that is, the proportion of records with the representative value of a trait to the total number of records for the trait of the species in a database. For *PH_max_*, we used the number of records with *PH* values. For the threshold values, we used 0.5, 0.6, 0.7, 0.8, 0.9, and 1.0, for *LS* and *GF* and 1, 2, 3, and 4 for *PH_max_*. The minimum threshold values (0.5 for *LS* and *GF*, and 1; *PH_max_*) imply no removal of the species. For each threshold value, trait, and database, we removed species with criterion values lower than the threshold and calculated the reliability index and corresponding novelty.

## RESULTS

### Number of species in each database

The number of species for each trait in each database is shown in Figure 1. The number of species obtained for *LS* was 131,059, 85,042, 47,137, 166,855, and 30,269 for the GBIF, TRY, BIEN, the Rice database, and eFloras, respectively (240,449 species for all databases). For *GF*, 149,067, 158,384, 93,329, 138,580, and 30,862 species were obtained from the GBIF, TRY, BIEN, Rice database, and eFloras, respectively (253,248 species for all databases). For *PH_max_*, 125,053, 31,397, 7896, and 30,774 species were obtained from the GBIF, TRY, BIEN, Rice database, and eFloras, respectively (148,601 species for all databases). The databases with the largest number of species were the rice database for *LS*, the TRY database for *GF*, and the GBIF database for *PH_max_*.

**FIGURE 1.**
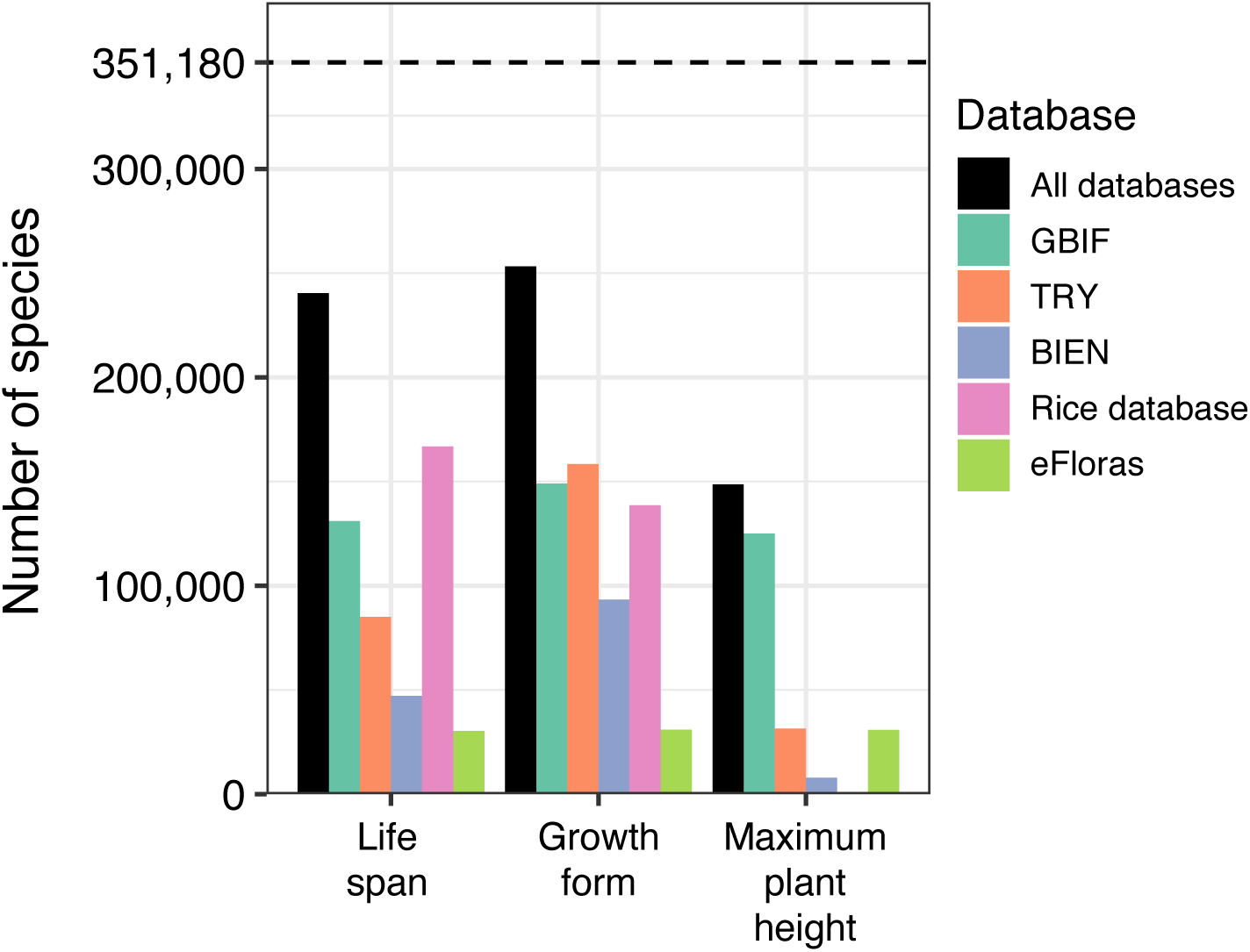
The number of species with trait values for each database and each trait. The dotted line indicates the total number of described vascular plant species (Freiberg et al., 2020).

### Reliability of plant traits in each database

For *LS* and *GF*, all databases contained proportion of species with congruent trait values with eFloras (i.e., reliability index for *LS* and *GF*) of 0.95 or higher, regardless of the threshold of criteria for removal of species with suspicious representative values (Figure 2A and B). However, for *GF*, the reliability index for GBIF was slightly lower than those of TRY and BIEN, regardless of the species removal threshold.

**FIGURE 2.**
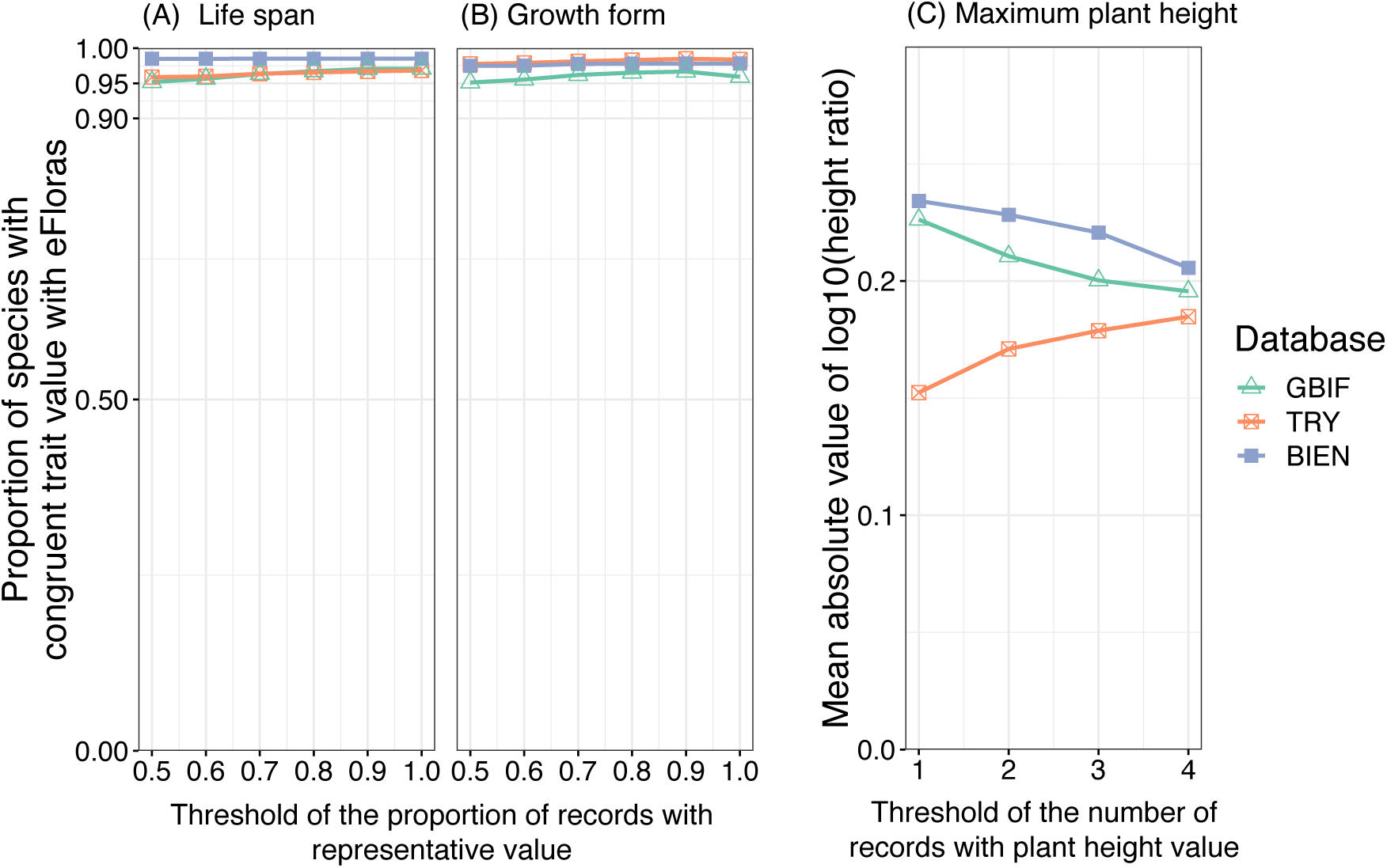
Changes in the reliability of (A) lifespan, (B) growth form, and (C) maximum plant height in each database along with the threshold of criteria for removal of species with suspicious values. The x-axes represent the threshold criteria used for the removal of species. Species with criterion values below the threshold were removed (i.e., higher x-axis values indicate the removal of more species). The minimum threshold values (0.5 for lifespan and growth form, and 1 for plant height) represent cases where no species were removed. The y-axis indicates the reliability index for each trait in each database. Points represent calculated values. Higher y-axis values for lifespan and growth form represent higher reliability. Lower y-axis values for maximum plant height represent higher reliability.

For *PH_max_*, the mean absolute value of the log10 *height ratio* of GBIF was lower than that of BIEN, that is, it had higher reliability than BIEN, regardless of the species removal threshold; however, TRY exhibited the highest reliability (Figure 2C).

### Novelty of plant traits in GBIF

We identified species with trait values obtained from GBIF that could not be obtained from other databases (Figure 3). Without removing species with suspicious representative values, the number of species solely obtained from GBIF was 37,422 for *LS*, 40,942 for *GF* and 93,129 for *PH_max_* (black parts with the minimum threshold values in Figure 3). When adding the number of species obtained solely from GBIF to the number of species obtained from the other four databases, the percentage of species for which trait values could be obtained for all described vascular plants increased from 58% to 68%, 60% to 72%, and 16% to 42% for *LS*, *GF*, and *PH_max_*, respectively. Furthermore, when the maximum threshold values for removing species were used in combination with suspicious trait values, large decreases in the number of species obtained solely from GBIF were not observed for *LS* or *GF* whereas a notable decrease was observed for *PH* when the maximum threshold value was used (from 93,129 to 57,054; Figure 3).

**FIGURE 3.**
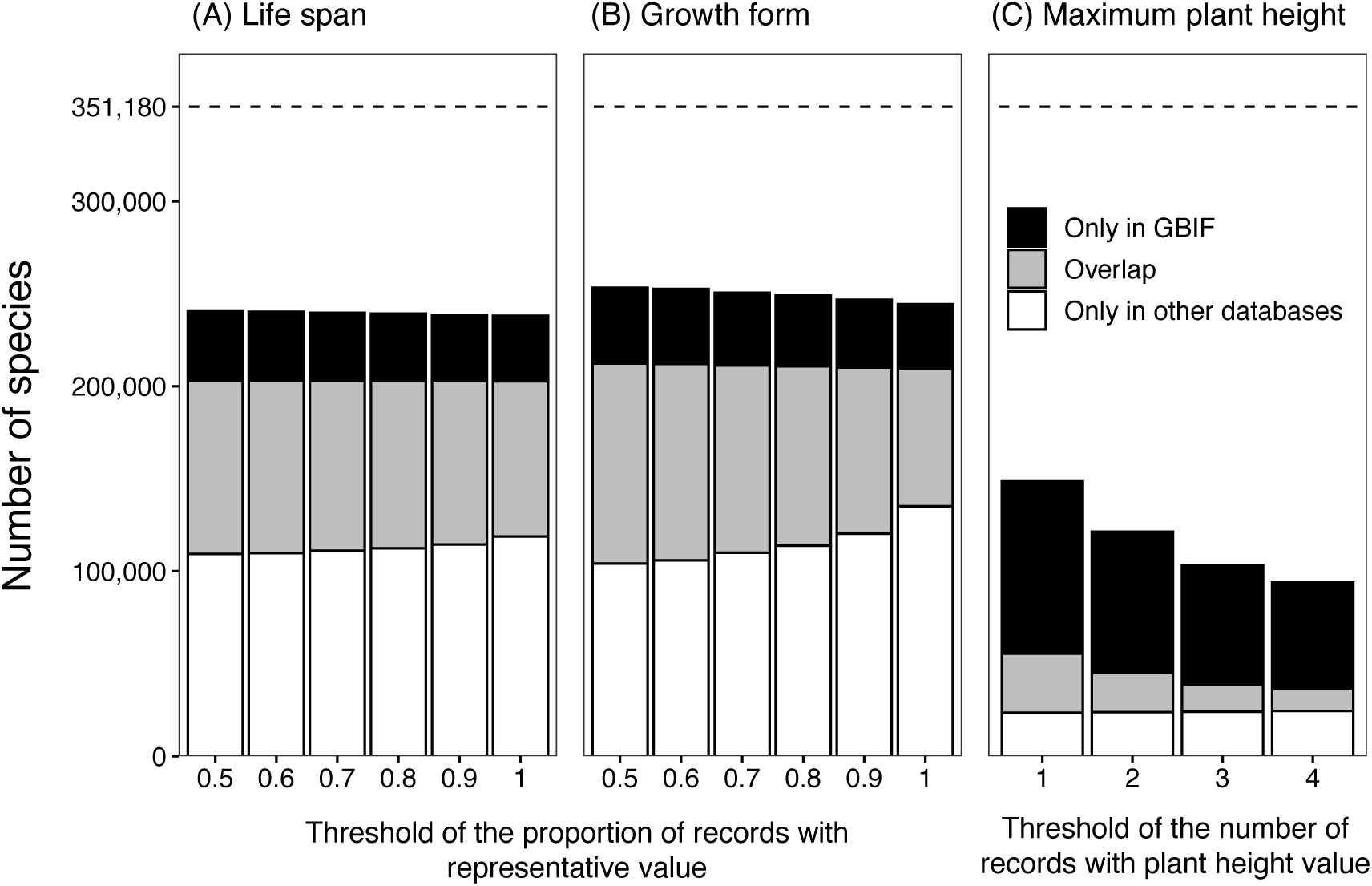
Changes in the number of species with (A) lifespan, (B) growth form, and (C) maximum plant height data obtained exclusively from GBIF, from other databases, or from both, following the removal of species with potentially incorrect values. The x-axis follows the same criteria as in Figure 2. Labels: *Only in GBIF*: species unique to GBIF; *Overlap*: species found in both GBIF and other databases; *Only in other databases*: species unique to the other databases. The total number of species in these three parts (= *Only in GBIF* + *Overlap* + *Only in other databases*) at the minimum threshold values are the same as the number of species of *All databases* in Figure 1. The dotted line represents the total number of described vascular plant species (Freiberg et al., 2020).

## DISCUSSION

### GBIF digital text as a new source for plant traits

Data on *LS*, *GF*, and *PH_max_* included in GBIF digital text had a reliability nearly similar to that of the common databases for plant traits (TRY or BIEN), even without the removal of species with suspicious representative values. Therefore, removing such species may not be necessary when using the *LS*, *GF*, and *PH_max_* of GBIF digital texts. Furthermore, *LS*, *GF*, and *PH_max_* data for many species are not included in the common databases for plant traits. Particularly, the number of species in *PH_max_* could be increased by approximately 2.7 times by adding data from the GBIF digital text. Therefore, we believe that the GBIF digital text is a useful and novel source of plant trait information. Plant trait information from GBIF’s digital text is anticipated to mitigate the severe Raunkiæran Shortfall (lack of trait information) and alleviate obstacles to plant trait research. However, the reliability of *GF* for GBIF was slightly lower than that of TRY and BIEN; therefore, caution may be necessary when using this database. This may be attributable to the presence of undiscovered errors in text mining (conversion, removal, and extraction of words) specific to *GF* in GBIF’s digital text. Furthermore, we used only North American and Chinese species to assess the reliability of plant traits. Therefore, if the reliability of the GBIF, TRY, and BIEN plant traits differs between these regions, the reliability of plant traits worldwide may not be accurately measured. Further studies are warranted to clarify this concern.

In addition to obtaining data from databases, other methods for obtaining missing trait information include manual literature research for each species or measurements from living individuals or specimens. However, these methods require considerable investment of time and effort. Therefore, obtaining considerable new trait data from the GBIF’s digital texts marks a significant milestone.

Although we used four databases (TRY, BIEN, eFloras, and the Rice database) as common databases for plant traits to compare *LS*, *GF*, and *PH_max_* with the GBIF digital text, several other databases include *LS*, *GF*, and/or *PH* information (Plant DNA C-values Database (https://cvalues.science.kew.org), USDA PLANTS Database (https://plants.usda.gov/adv_search.html), Jepson eFlora (https://ucjeps.berkeley.edu/eflora/), Angiosperm Phylogeny Website (http://www.mobot.org/MOBOT/research/Apweb/), AusTraits (Falster et al., 2021), etc.). These four databases were selected as they are the most common ones and enable ease of extraction, bearing in mind the balance between effort invested and research quality. Therefore, the novelty of GBIF’s digital text demonstrated in this study is only relevant when compared with the other four databases. If the number of databases and literature used increases, the novelty of GBIF’s digital text may decrease.

Furthermore, each GBIF record (occurrence) may include location information (coordinates and locations). Therefore, combining the trait and location information in each record may allow for the examination of geographic variation in traits. For instance, examining geographical variations in *PH* within species may be beneficial.

In this study, we extracted *LS*, *GF*, and *PH_max_* information from the GBIF and common databases for plant traits (TRY, BIEN, Rice database, and eFloras), and organized them at the species level. Consequently, a data table with the largest number of species for these three traits was created (see Data Availability Statement). This data table can be used for future research on plant traits. To the best of our knowledge, this study is the first to extract plant trait information from GBIF’s digital textual data of the GBIF. Although plant trait information from four databases other than GBIF has been extracted and used in previous studies (e.g. Thomson et al., 2011; Diaz et al., 2016; Kling and Ackerly, 2021; Poppenwimer et al., 2023), it appears that consolidating the data into one data table offers a convenient approach.

### Reasons why trait information in GBIF’s digital text has not been used

Although GBIF’s digital text contains plant trait information for a large number of species, it has rarely been extracted and used. Two possible reasons may explain this. First, GBIF is primarily recognized as a database for information on species distribution (König et al., 2019), making it likely that researchers overlook trait information in the textual data. Second, the trait values included in the digital text are presented as sentences rather than words and phrases (Table 2). Therefore, trait information is not easily accessible, requiring efforts such as eliminating phrases that are not related to traits and extracting trait information from sentences.

### Unextracted trait information that can be extracted from GBIF

In this study, we extracted only *LS*, *GF*, and *PH_max_* from GBIF digital text; however, we observed that GBIF digital text also contains information on leaf habit (evergreen or deciduous), climbing habit (climber or erect), epiphytic habit (epiphyte or geophyte), and flower color (yellow, white, pink, etc.) (Table 3). These traits are important for understanding plant growth, phenology, and pollination (Rowe and Speck, 2005; Chittka and Raine, 2006; Rausher, 2008; van Ommen Kloeke et al., 2012; Gianoli, 2015; Zotz, 2016). Therefore, it is important to extract these traits. However, in this study, we did not extract these traits because of labor limitations; therefore, we hope future studies will address this gap. Furthermore, recent studies have attempted to extract plant trait information from taxonomic literature on plants (Coleman et al., 2023; Folk et al., 2024). By applying these methods, it may be possible to extract data on a broader range of species and types of traits from the GBIF digital text than that achieved in this study.

**TABLE 3.**
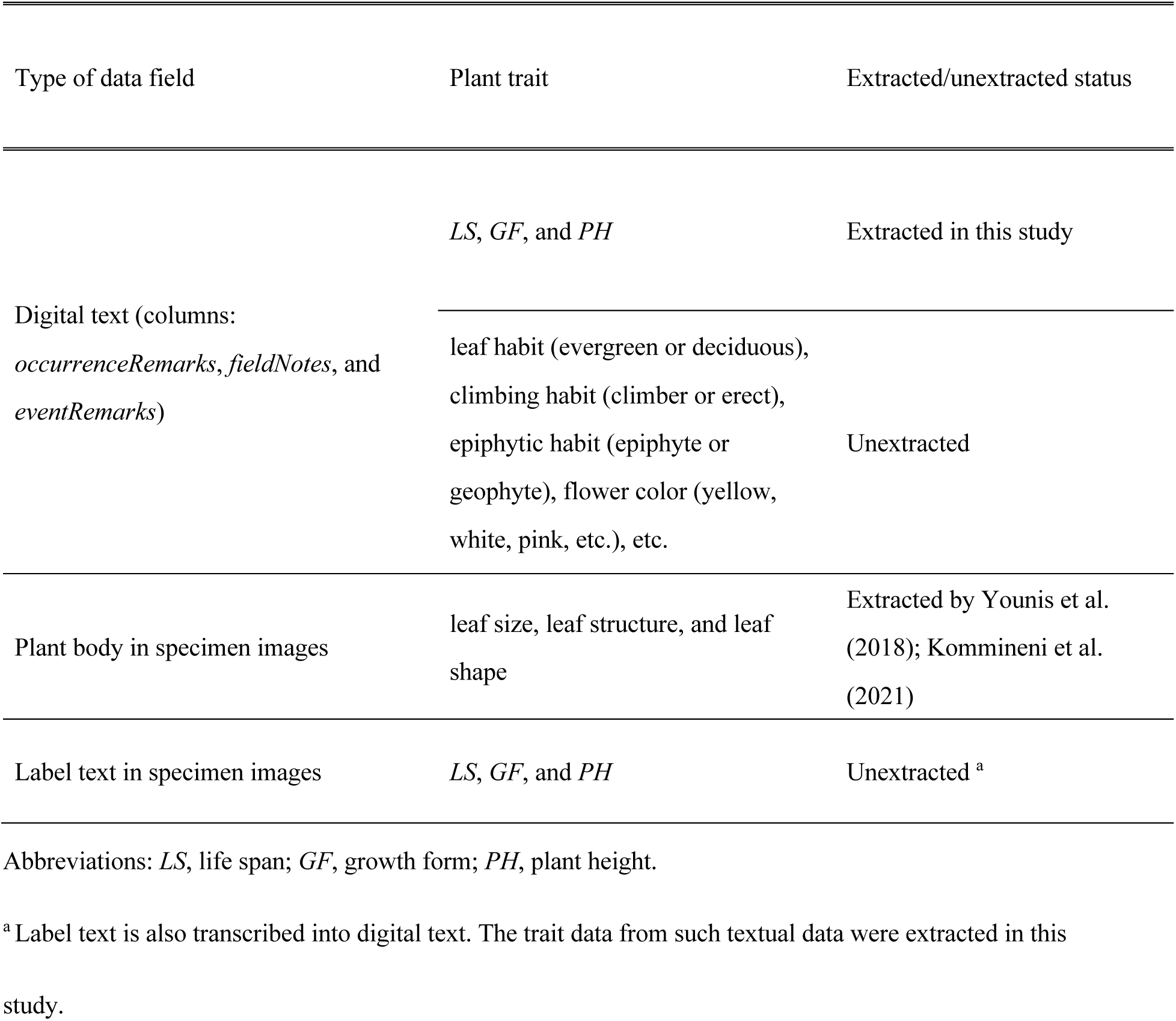
Utilization status of plant trait information included in GBIF.

Furthermore, GBIF contains plant trait information not only in its digital text but also in specimen image data (Table 3). Specifically, plant trait information can be obtained from each plant part and labeled part of the specimen image. Plant trait information (leaf size and shape) from the plant part has previously been obtained (Younis et al., 2018; Kommineni et al., 2021). Some of the labeled parts have already been transcribed into the GBIF digital text data; therefore, they were included in the data extracted in this study (Table 3). However, not all labels have been transcribed into digital text, leaving parts of the data unextracted (see Table 3). Numerous attempts have been made to transcribe label text into digital text (Greenberg and Klas, 2008; Kirchhoff et al., 2018; Nelson and Ellis, 2019; Owen et al., 2020; Walton et al., 2020; Albani Rocchetti et al., 2021; Weaver et al., 2023; Folk et al., 2024). We anticipate that some of the GBIF label images that have not yet been transcribed into digital text will be digitized in the future.

Furthermore, as GBIF includes records of organisms other than plants, it is possible that of the digital text, specimen body images, and specimen label images in their records also contain useful trait information.

### Lack of plant trait information that cannot be resolved with GBIF

As described in the Materials and Methods section, the main functional traits of plants are divided into size-related traits (e.g., *LS*, *GF*, *PH_max_*, and leaf area) and leaf economy-related traits (e.g., leaf mass per area and leaf nitrogen content) (Diaz et al., 2016). Of these, size-related traits can be extracted from GBIF digital text or specimen images (plant bodies or labels); however, leaf economy-related traits are unlikely to be retrievable from GBIF. To supplement the leaf economy-related traits of species that are not included in the common plant trait database, acquiring them from sources other than the GBIF is necessary. Possible methods include extraction from various studies and destructive sampling of specimens or live individuals (Heberling, 2022).

## CONCLUSIONS

The digital text data from the GBIF has rarely been used as a source of plant trait information. However, it contains plant trait information for species that are not available in common databases; furthermore, it demonstrated a reliability comparable to that the common databases. Therefore, GBIF’s digital texts are a promising new source of plant trait information that may alleviate the lack of plant trait information and contribute to advancing plant trait research. Furthermore, by extracting and organizing traits from five databases, including GBIF, we created a data table containing the largest number of species for *LS*, *GF*, and *PH_max_*. Additionally, GBIF contains additional trait information that has not been extracted in this study, highlighting an important area for future research.

## Supporting information

Appendix S1. Supplementary information on methods for extracting trait values.

## ACKNOWLEDGMENTS

This study was supported in part by the Environment Research and Technology Development Fund (JPMEERF23S12103) of the Environmental Restoration and Conservation Agency, provided by the Ministry of the Environment of Japan.

## AUTHOR CONTRIBUTIONS

T.K. conceptualized the objectives and methodology of this study. M. O. refined the methodology. T. K. and M. O. conducted the analyses and wrote the manuscript. All authors approved the final version of the manuscript.

## DATA AVAILABILITY STATEMENT

The data associated with this study are available on Zenodo (DOI: doi.org/10.5281/zenodo.15030184) and include: 1) a raw table presenting the occurrence data on vascular plants downloaded from GBIF, containing the four columns used in this study as well as latitude and longitude data; 2) expressions of *PH* values extracted from each GBIF record; 3) a species-level dataset of *LS*, *GF*, and *PH_max_* for each database.

Additional Supporting Information can be found in the Supporting Information section at the end of this article.

AAppendix S1. Supplementary information on methods for extracting trait values.

## LITERATURE CITED

Albani Rocchetti, G., C. G. Armstrong, T. Abeli, S. Orsenigo, C. Jasper, S. Joly, A. Bruneau, et al. 2021. Reversing extinction trends: new uses of (old) herbarium specimens to accelerate conservation action on threatened species. New Phytologist 230: 433–450.

Bernal, R., B. Martínez, and M. J. Sanín. 2018. The World’s Tallest Palms. Palms 62.

Boonman, C. C. F., A. Benítez-López, A. M. Schipper, W. Thuiller, M. Anand, B. E. L. Cerabolini, J. H. C. Cornelissen, et al. 2020. Assessing the reliability of predicted plant trait distributions at the global scale. Global Ecology and Biogeography 29: 1034–1051.

Bruelheide, H., J. Dengler, O. Purschke, J. Lenoir, B. Jimenez-Alfaro, S. M. Hennekens, Z. Botta-Dukat, et al. 2018. Global trait-environment relationships of plant communities. Nature Ecology & Evolution 2: 1906–1917.

Butler, E. E., A. Datta, H. Flores-Moreno, M. Chen, K. R. Wythers, F. Fazayeli, A. Banerjee, et al. 2017. Mapping local and global variability in plant trait distributions. Proceedings of the National Academy of Sciences 114: E10937–E10946.

Chen, S. C., R. Tamme, F. J. Thomson, and A. T. Moles. 2019. Seeds tend to disperse further in the tropics. Ecology Letters 22: 954–961.

Chittka, L., and N. E. Raine. 2006. Recognition of flowers by pollinators. Current Opinion in Plant Biology 9: 428–435.

Coleman, D., R. V. Gallagher, D. Falster, H. Sauquet, and E. Wenk. 2023. A workflow to create trait databases from collections of textual taxonomic descriptions. Ecological Informatics 78.

Cornwell, W. K., W. D. Pearse, R. L. Dalrymple, and A. E. Zanne. 2019. What we (don’t) know about global plant diversity. Ecography 42: 1819–1831.

Diaz, S., J. Kattge, J. H. Cornelissen, I. J. Wright, S. Lavorel, S. Dray, B. Reu, et al. 2016. The global spectrum of plant form and function. Nature 529: 167–171.

eFloras. 2008. Published on the Internet http://www.efloras.org [accessed 26 August 2023]. Missouri Botanical Garden, St. Louis, MO & Harvard University Herbaria Cambridge, MA.

Falster, D., R. Gallagher, E. H. Wenk, I. J. Wright, D. Indiarto, S. C. Andrew, C. Baxter, et al. 2021. AusTraits, a curated plant trait database for the Australian flora. Scientific Data 8: 254.

Foden, W. B., S. H. Butchart, S. N. Stuart, J.-C. Vié, H. R. Akçakaya, A. Angulo, L. M. DeVantier, et al. 2013. Identifying the world’s most climate change vulnerable species: a systematic trait-based assessment of all birds, amphibians and corals. PLoS One 8: e65427.

Folk, R. A., R. P. Guralnick, and R. T. LaFrance. 2024. FloraTraiter: Automated parsing of traits from descriptive biodiversity literature. Applications in Plant Sciences: e11563.

Freiberg, M., M. Winter, A. Gentile, A. Zizka, A. N. Muellner-Riehl, A. Weigelt, and C. Wirth. 2020. LCVP, The Leipzig catalogue of vascular plants, a new taxonomic reference list for all known vascular plants. Scientific Data 7: 1–7.

GBIF Secretariat. 2021. GBIF Science Review 2020.

GBIF: The Global Biodiversity Information Facility. 2025. What is GBIF? https://www.gbif.org/what-is-gbif [accessed 8 March 2025].

GBIF.org. 2021. GBIF Occurrence. 10.15468/dl.qpuj6c [downloaded 13 September 2021].

Gianoli, E. 2015. The behavioural ecology of climbing plants. AoB Plants 7: plv013.

Greenberg, J., and W. Klas. 2008. Metadata for semantic and social applications DC-2008 Berlin-Proceedings of the 8. International conference on Dublin Core and Metadata Applications. Universitätsverlag Göttingen.

Grubbs, F. E. 1969. Procedures for detecting outlying observations in samples. Technometrics 11: 1–21.

Hawkins, B. A., M. Á. Rodríguez, and S. G. Weller. 2011. Global angiosperm family richness revisited: linking ecology and evolution to climate. Journal of Biogeography 38: 1253–1266.

Heberling, J. M. 2022. Herbaria as Big Data Sources of Plant Traits. International Journal of Plant Sciences 183: 87–118.

Hoffmann, A. A., and C. M. Sgrò. 2011. Climate change and evolutionary adaptation. Nature 470: 479–485.

Hortal, J., F. de Bello, J. A. F. Diniz-Filho, T. M. Lewinsohn, J. M. Lobo, and R. J. Ladle. 2015. Seven shortfalls that beset large-scale knowledge of biodiversity. Annual Review of Ecology, Evolution, and Systematics 46: 523–549.

Kattge, J., G. Bönisch, S. Díaz, S. Lavorel, I. C. Prentice, P. Leadley, S. Tautenhahn, et al. 2020. TRY plant trait database–enhanced coverage and open access. Global Change Biology 26: 119–188.

Kattge, J., S. DÍAz, S. Lavorel, I. C. Prentice, P. Leadley, G. BÖNisch, E. Garnier, et al. 2011. TRY - a global database of plant traits. Global Change Biology 17: 2905–2935.

Kelly, R., K. Healy, M. Anand, M. E. Baudraz, M. Bahn, B. E. Cerabolini, J. H. Cornelissen, et al. 2021. Climatic and evolutionary contexts are required to infer plant life history strategies from functional traits at a global scale. Ecology Letters 24: 970–983.

Kirchhoff, A., U. Bügel, E. Santamaria, F. Reimeier, D. Röpert, A. Tebbje, A. Güntsch, et al. 2018. Toward a service-based workflow for automated information extraction from herbarium specimens. Database 2018.

Kling, M. M., and D. D. Ackerly. 2021. Global wind patterns shape genetic differentiation, asymmetric gene flow, and genetic diversity in trees. Proceedings of the National Academy of Sciences 118: e2017317118.

Kommineni, V. K., S. Tautenhahn, P. Baddam, J. Gaikwad, B. Wieczorek, A. Triki, and J. Kattge. 2021. Comprehensive leaf size traits dataset for seven plant species from digitised herbarium specimen images covering more than two centuries. Biodiversity Data Journal 9.

König, C., P. Weigelt, J. Schrader, A. Taylor, J. Kattge, and H. Kreft. 2019. Biodiversity data integration—the significance of data resolution and domain. PLoS Biology 17: e3000183.

König, C., P. Weigelt, A. Taylor, A. Stein, W. Dawson, F. Essl, J. Pergl, et al. 2021. Source pools and disharmony of the world’s island floras. Ecography 44: 44–55.

Lamanna, C., B. Blonder, C. Violle, N. J. Kraft, B. Sandel, I. Simova, J. C. Donoghue, 2nd, et al. 2014. Functional trait space and the latitudinal diversity gradient. Proceedings of the National Academy of Sciences, USA 111: 13745–13750.

Maitner, B. S., R. Gallagher, J.-C. Svenning, M. Tietje, E. H. Wenk, and W. L. Eiserhardt. 2022. Socioeconomics and biogeography jointly drive geographic biases in our knowledge of plant traits: a global assessment of the Raunkiaerian shortfall in plants. bioRxiv.

Maitner, B. S., B. Boyle, N. Casler, R. Condit, J. Donoghue II, S. M. Durán, D. Guaderrama, et al. 2018. The bien r package: A tool to access the Botanical Information and Ecology Network (BIEN) database. Methods in Ecology and Evolution 9: 373–379.

Moles, A. T., and M. R. Leishman. 2008. The seedling as part of a plant’s life history strategy. In M. A. Leck, V. T. Parker, and R. L. Simpson [eds.], Seedling Ecology and Evolution. Cambridge University Press, Cambridge.

Moles, A. T., D. I. Warton, L. Warman, N. G. Swenson, S. W. Laffan, A. E. Zanne, A. Pitman, et al. 2009. Global patterns in plant height. Journal of Ecology 97: 923–932.

Morin, X., and I. Chuine. 2006. Niche breadth, competitive strength and range size of tree species: a trade-off based framework to understand species distribution. Ecology Letters 9: 185–195.

Nelson, G., and S. Ellis. 2019. The history and impact of digitization and digital data mobilization on biodiversity research. Philosophical Transactions of the Royal Society B 374: 20170391.

Ordoñez, J. C., P. M. Van Bodegom, J. P. M. Witte, I. J. Wright, P. B. Reich, and R. Aerts. 2009. A global study of relationships between leaf traits, climate and soil measures of nutrient fertility. Global Ecology and Biogeography 18: 137–149.

Owen, D., L. Livermore, Q. Groom, A. Hardisty, T. Leegwater, M. van Walsum, N. Wijkamp, and I. Spasic. 2020. Towards a scientific workflow featuring Natural Language Processing for the digitisation of natural history collections. Research Ideas and Outcomes 6.

Perez, T. M., O. Valverde-Barrantes, C. Bravo, T. C. Taylor, B. Fadrique, J. A. Hogan, C. J. Pardo, et al. 2019. Botanic gardens are an untapped resource for studying the functional ecology of tropical plants. Philosophical Transactions of the Royal Society B: Biological Sciences 374: 20170390.

Pierce, S., D. Negreiros, B. E. L. Cerabolini, J. Kattge, S. Díaz, M. Kleyer, B. Shipley, et al. 2017. A global method for calculating plant CSR ecological strategies applied across biomes world- wide. Functional Ecology 31: 444–457.

Poppenwimer, T., I. Mayrose, and N. DeMalach. 2023. Revising the global biogeography of annual and perennial plants. Nature 624: 109–114.

R-Core-Team. 2021. R: A language and environment for statistical computing. website: https://www.R-project.org/.

Rausher, M. D. 2008. Evolutionary transitions in floral color. International Journal of Plant Sciences 169: 7–21.

Rice, A., P. Smarda, M. Novosolov, M. Drori, L. Glick, N. Sabath, S. Meiri, et al. 2019. The global biogeography of polyploid plants. Nature Ecology & Evolution 3: 265–273.

Rowe, N., and T. Speck. 2005. Plant growth forms: an ecological and evolutionary perspective. New Phytologist 166: 61–72.

Sandel, B., A. G. Gutiérrez, P. B. Reich, F. Schrodt, J. Dickie, and J. Kattge. 2015. Estimating the missing species bias in plant trait measurements. Journal of Vegetation Science 26: 828–838.

Shenkin, A., C. J. Chandler, D. S. Boyd, T. Jackson, M. Disney, N. Majalap, R. Nilus, et al. 2019. The world’s tallest tropical tree in three dimensions. Frontiers in Forests and Global Change 2: 32.

Shiklomanov, A. N., E. M. Cowdery, M. Bahn, C. Byun, S. Jansen, K. Kramer, V. Minden, et al. 2020. Does the leaf economic spectrum hold within plant functional types? A Bayesian multivariate trait meta-analysis. Ecological Applications 30: e02064.

Sillett, S. C., R. D. Kramer, R. Van Pelt, A. L. Carroll, J. Campbell-Spickler, and M. E. Antoine. 2021. Comparative development of the four tallest conifer species. Forest Ecology and Management 480: 118688.

Šímová, I., C. Violle, J. C. Svenning, J. Kattge, K. Engemann, B. Sandel, R. K. Peet, et al. 2018. Spatial patterns and climate relationships of major plant traits in the New World differ between woody and herbaceous species. Journal of Biogeography 45: 895–916.

Smith, S. A., and M. J. Donoghue. 2008. Rates of molecular evolution are linked to life history in flowering plants. Science 322: 86–89.

Smith, S. A., and J. M. Beaulieu. 2009. Life history influences rates of climatic niche evolution in flowering plants. Proceedings of the Royal Society of London B: Biological Sciences 276: 4345–4352.

Soltis, D. E., M. E. Mort, M. Latvis, E. V. Mavrodiev, B. C. O’Meara, P. S. Soltis, J. G. Burleigh, and R. Rubio de Casas. 2013. Phylogenetic relationships and character evolution analysis of Saxifragales using a supermatrix approach. American Journal of Botany 100: 916–929.

Stebbins, G. L. 1971. Chromosomal evolution in higher plants. Edward Arnold Ltd., London.

Swenson, N. G., and M. D. Weiser. 2010. Plant geography upon the basis of functional traits: an example from eastern North American trees. Ecology 91: 2234–2241.

Thomson, F. J., A. T. Moles, T. D. Auld, and R. T. Kingsford. 2011. Seed dispersal distance is more strongly correlated with plant height than with seed mass. Journal of Ecology 99: 1299–1307.

Valladares, F., E. Gianoli, and J. M. Gómez. 2007. Ecological limits to plant phenotypic plasticity. New Phytologist 176: 749–763.

Vallicrosa, H., J. Sardans, J. Maspons, and J. Peñuelas. 2022. Global distribution and drivers of forest biome foliar nitrogen to phosphorus ratios (N:P). Global Ecology and Biogeography 31: 861–871.

van Ommen Kloeke, A., J. Douma, J. C. Ordonez, P. B. Reich, and P. Van Bodegom. 2012. Global quantification of contrasting leaf life span strategies for deciduous and evergreen species in response to environmental conditions. Global Ecology and Biogeography 21: 224–235.

Violle, C., M. L. Navas, D. Vile, E. Kazakou, C. Fortunel, I. Hummel, and E. Garnier. 2007. Let the concept of trait be functional! Oikos 116: 882–892.

Walton, S., L. Livermore, O. Bánki, R. Cubey, R. Drinkwater, M. Englund, C. Goble, et al. 2020. Landscape analysis for the specimen data refinery. Research Ideas and Outcomes 6: e57602.

Wang, K. L., P. R. Deng, Z. Yao, J. Y. Dong, Z. He, P. Yang, and Y. B. Liu. 2023. Biogeographic patterns of polyploid species for the angiosperm flora in China. Journal of Systematics and Evolution 61: 776–789.

Weaver, W. N., B. R. Ruhfel, K. J. Lough, and S. A. Smith. 2023. Herbarium specimen label transcription reimagined with large language models: Capabilities, productivity, and risks. American Journal of Botany 110: e16256.

Wee, M. S., L. Matia-Merino, S. M. Carnachan, I. M. Sims, and K. K. Goh. 2014. Structure of a shear- thickening polysaccharide extracted from the New Zealand black tree fern, Cyathea medullaris. International Journal of Biological Macromolecules 70: 86–91.

Westoby, M., D. S. Falster, A. T. Moles, P. A. Vesk, and I. J. Wright. 2002. Plant ecological strategies: some leading dimensions of variation between species. Annual Review of Ecology and Systematics: 125–159.

Wright, I. J., P. B. Reich, M. Westoby, D. D. Ackerly, Z. Baruch, F. Bongers, J. Cavender-Bares, et al. 2004. The worldwide leaf economics spectrum. Nature 428: 821–827.

Younis, S., C. Weiland, R. Hoehndorf, S. Dressler, T. Hickler, B. Seeger, and M. Schmidt. 2018. Taxon and trait recognition from digitized herbarium specimens using deep convolutional neural networks. Botany Letters 165: 377–383.

Zanne, A. E., D. C. Tank, W. K. Cornwell, J. M. Eastman, S. A. Smith, R. G. FitzJohn, D. J. McGlinn, et al. 2014. Three keys to the radiation of angiosperms into freezing environments. Nature 506: 89–92.

Zotz, G. 2016. Plants on plants-the biology of vascular epiphytes. Springer.

