## Appendix S1. Supplementary information on methods for extracting trait values. for "Extracting and evaluating plant trait information from digital text in the Global Biodiversity Information Facility"

**Appendix S1. Supplementary information on methods used for extracting trait values**

**Table S1**. Sources of trait information for each trait and database.

| Database | Trait | Source(s) of trait information |
|---|---|---|
| GBIF | *LS*, *GF*, and *PH* | *occurrenceRemarks* column: notes on the occurrence |
| | | *fieldNotes* column: field note |
| | | *eventRemarks* column: notes on the event |
| TRY | *LS* and *GF* | *TraitName* column: name of trait |
| | | *DataName* column: name of specific trait or context information |
| | | *OrigValueStr* column: original value as text string |
| | *PH* | *TraitName* column: name of trait |
| | | *OrigValueStr* column: original value as text string |
| | | *StdValue* column: standardized value |
| | | *UnitName* column: unit name |
| BIEN | *LS*, *GF*, and *PH* | *trait_name* column |
| | | *trait_value* column |
| Rice's database | *LS* | *Life_cycle* column |
| | *GF* | *Growth_form* column |
| eFloras | *LS*, *GF*, and *PH* | First sentence of the description of each taxon of each flora |

Abbreviations: *LS*, life span; *GF*, growth form; *PH*, plant height.

**Table S2**. List of expressions excluded prior to the extraction of values for life span and growth form.

| Category | Expression | Note |
|---|---|---|
| Precipitation | *annual rainfall* | |
| | *annual precipitation* | |
| Branch form | *一年生枝* | Current year branch (in Chinese). |
| | *二年生枝* | Second year branch (in Chinese). |
| Woodiness of the plant part | *woody base* | |
| | *woody-base* | |
| | *woody root* | |
| | *woody-root* | |
| | *woody at* | |
| Abbreviation for herbarium | *ex herb.* | |
| | *ex. herb.* | |
| | *ex-herb.* | |
| | *ex- herb.* | |
| | *ex - herb.* | |
| Living environment | Combinations of | e.g., *on a high shrub*, *among the trees*, etc. |
| | [*on*, *of*, *among*, *under*] | |
| | [*a*, *the*] | |
| | [*low*, *high*] | |
| | [*tree*, *shrub*] | |
| Farming practice | *bienal modal* | e.g., *campo sujo com queima bienal modal* |

**Table S3**. Expressions representing life span (*LS*) and growth form (*GF*) used to extract trait values from the GBIF text. Italics denote the expressions used for the search without alterations. To include expressions of *LS* and *GF* with the prefix *sub-* (e.g., subshrub) and suffix *-let* (e.g., shrublet), these prefixes and suffixes were excluded prior to the extraction of trait values.

| Expressions representing the class of the trait | Resolved class |
|---|---|
| *anual* | Annual |
| *annual* | Annual |
| *annuelle* | Annual |
| *一年* | Annual |
| *biennial* | Annual |
| *bienal* | Annual |
| *二年* | Annual |
| *perennial* | Perennial |
| *perenne* | Perennial |
| *多年* | Perennial |
| *erva* | Herb |
| *forb* | Herb |
| *gras* | Herb |
| *grass* | Herb |
| *herb* | Herb |
| *herbacea* | Herb |
| *herbaceo* | Herb |
| *herbaceous* | Herb |
| *herbe* | Herb |
| *hierba* | Herb |
| *sedge* | Herb |
| *weed* | Herb |
| *性状:h* | Herb |
| *草本* | Herb |
| *草質藤本* | Herb |

**Table S3**. Continued.

| Expressions representing the class of the trait | Resolved class |
|---|---|
| *abusto* | Wood |
| *arbiusto* | Wood |
| *arbol* | Wood |
| *arbolito* | Wood |
| *arborea* | Wood |
| *arboreo* | Wood |
| *arborescente* | Wood |
| *arbre* | Wood |
| *arbrisseau* | Wood |
| *arbusot* | Wood |
| *arbust* | Wood |
| *arbuste* | Wood |
| *arbustiva* | Wood |
| *arbustivo* | Wood |
| *arbusto* | Wood |
| *arvore* | Wood |
| *arvoreta* | Wood |
| *arvoreto* | Wood |
| *avore* | Wood |
| *avoreto* | Wood |
| *bark* | Wood |
| *baum* | Wood |
| *bush* | Wood |
| *liana* | Wood |
| *mallee* | Wood |
| *shrub* | Wood |
| *shurb* | Wood |
| *suffrutex* | Wood |
| *tree* | Wood |
| *woody* | Wood |
| *喬木* | Wood |
| *性狀:sh* | Wood |
| *性狀:t* | Wood |
| *木質藤本* | Wood |
| *木本* | Wood |
| *灌木* | Wood |

**Table S4**. Expressions representing life span (*LS*) and growth form (*GF*) used to extract trait values from eFloras. Italics denote the expressions used for the search without alteration. To include expressions of *LS* and *GF* with the prefix *sub-* (e.g., subshrub) and suffix *-let* (e.g., shrublet), these prefixes and suffixes were excluded prior to the extraction of trait values.

| Expressions representing the class of the trait | Resolve class |
|---|---|
| *annual* | Annual |
| *biennial* | Annual |
| *perennial* | Perennial |
| *herb* | Herb |
| *vine herbaceous* | Herb |
| *vine, herbaceous* | Herb |
| *climber, herbaceous* | Herb |
| *herbaceous climber* | Herb |
| *herbaceous vine* | Herb |
| *tree* | Wood |
| *shrub* | Wood |
| *liana* | Wood |
| *vine woody* | Wood |
| *vine, woody* | Wood |
| *climber woody* | Wood |
| *climber, woody* | Wood |
| *woody climber* | Wood |
| *woody vine* | Wood |

**Table S5**. Combination of values in the *TraitNames*, *DataName*, and *OrigValueStr* columns used to assign candidate values of life span (*LS*) and growth form (*GF*) for each record in TRY. For the matching method (a), values of three columns aligned in each row were matched, whereas all combinations of *TraitName* and *OrigValueStr* were considered for method (b). Italics denote the expressions used for the search. When assigning trait values, partial matches of expressions with cell contents were permitted, whereas expressions shown in bold italics were evaluated based on an exact match with the cell contents.

| Trait | Matching method | Corresponding values in columns | | | Resolved class |
|---|---|---|---|---|---|
| | | *TraitName* column | *DataName* column | *OrigValueStr* column | |
| *LS* | (a) | *Plant lifespan (longevity)* | Any | Numbers ≤ 2 | Annual |
| | | | Any | Numbers > 2 | Perennial |
| | | | Any | *poly-annuals* | Perennial |
| | | | *Plant phenology: Annual* | ***yes*** | Annual |
| | | | *Plant phenology: Biennial* | ***yes*** | Annual |
| | | | *Plant life span: annual* | ***yes*** | Annual |
| | | | *Plant phenology: Perennial* | ***yes*** | Perennial |
| | | | *Plant life span: perennial* | ***yes*** | Perennial |
| | (b) | *Plant lifespan (longevity)*, *Plant growth form*, *Plant life form (Raunkiaer life form)*, *Plant woodiness*, *Plant vegetative regeneration capacity*, *Plant functional type (PFT)* | Any | *annual* | Annual |
| | | | Any | ***ann*** | Annual |
| | | | Any | *biannual* | Annual |
| | | | Any | *biennial* | Annual |
| | | | Any | *biennual* | Annual |
| | | | Any | *bieenial* | Annual |
| | | | Any | *bennial* | Annual |
| | | | Any | *perennial* [a] | Perennial |
| | | | Any | *pernnial* | Perennial |
| | | | Any | *pluriennial* | Perennial |
| | | | Any | ***pere*** | Perennial |
| | | | Any | ***1, 2, more*** | Perennial |

Note:

[a] When it appeared as *not perennial*, the occurrence was ignored.

**Table S5**. Continued.

| Trait | Matching method | Correspondent values in columns | | | Resolved class |
|---|---|---|---|---|---|
| | | *TraitName* column | *DataName* column | *OrigValueStr* column | |
| *GF* | (b) | *Plant lifespan (longevity)*, *Plant growth form*, *Plant life form (Raunkiaer life form)*, *Plant woodiness*, *Plant vegetative regeneration capacity*, *Plant functional type (PFT)* | Any | *herb* | Herb |
| | | | Any | ***h*** | Herb |
| | | | Any | *forb* | Herb |
| | | | Any | *herbaceous* | Herb |
| | | | Any | *gras* | Herb |
| | | | Any | *grass* | Herb |
| | | | Any | *graminoid* | Herb |
| | | | Any | *non-woody* | Herb |
| | | | Any | *non woody* | Herb |
| | | | Any | *false* | Herb |
| | | | Any | tree | Wood |
| | | | Any | ***t*** | Wood |
| | | | Any | ***w*** | Wood |
| | | | Any | ***s*** | Wood |
| | | | Any | *shrub* | Wood |
| | | | Any | *woody* [b] | Wood |
| | | | Any | *liana* | Wood |
| | | | Any | *palm* | Wood |
| | | | Any | *bamboo* | Wood |
| | | | Any | *mallee* | Wood |
| | | | Any | *true* | Wood |

Note:

[b] However, *non woody* and *non-woody* was classified as a herb.

**Table S6**. Combination of values in the *trait_name* and *trait_value* columns used to assign candidate values of life span (*LS*) and growth form (*GF*) for each record in BIEN. For matching method (a), all combinations of *trait_name* and *trait_value* were considered when assigning candidate trait values, whereas the values of the two columns aligned in each row were matched together for method (b). Italics denote the expressions used for the search. When assigning trait values, partial matches of expressions with cell contents were permitted.

| Trait | Matching method | Correspondent values in columns | | Resolved class |
|---|---|---|---|---|
| | | *trait_name* column | *trait_value* column | |
| *LS* | (a) | *longest whole plant longevity*, *maximum whole plant longevity* | Numbers ≤ 2 | Annual |
| | | | Numbers > 2 | Perennial |
| | (b) | *whole plant growth form* | *pere* | Perennial |
| *GF* | (b) | *whole plant growth form*, *whole plant growth form diversity*, *whole plant woodiness* | *herb* | Herb |
| | | | *forb* | Herb |
| | | | *herbaceous* | Herb |
| | | | *gras* | Herb |
| | | | *grass* | Herb |
| | | | *graminoid* | Herb |
| | | | *non-woody* | Herb |
| | | | *tree* | Wood |
| | | | *shrub* | Wood |
| | | | *woody* [a] | Wood |
| | | | *liana* | Wood |
| | | | *palm* | Wood |
| | | | *bamboo* | Wood |

Note: [a] *non-woody* plants are regarded as herbs.

**Table S7**. List of expressions representing approximations excluded prior the extraction of plant height values

| Expressions |
|---|
| *c* |
| *c.* |
| *ca* |
| *ca.* |
| *ca.,* |
| *c.a.* |
| *cerca* |
| *about* |
| *aproximadamente* |
| *aprox.* |
| *approx.* |
| *approximately* |
| *aproximada* |
| *約* |
| ± |

Note: In cases where *de* was placed between the expressions listed above and a number (e.g., *ca de 1*), it was excluded from the search.

**Table S8**. Patterns of expression that could potentially cause false detection of plant height values were removed from the records in GBIF and eFloras. When one of these expressions was placed immediately before or after the combination of number and unit (e.g. *10 m width*, *width 10m*, etc.), the combination of the number, unit, and expression was removed. The numbers included integers, decimals, fractions, and mixed fractions, and the units are listed in Table S9. The combination of numbers and units can be within this range. An asterisk following an expression indicates that all the words starting from the expression were used.

| Order of expression, number, and unit | Category of expression | Expression | Note |
|---|---|---|---|
| An expression is placed immediately **after** the combination of number and unit (e.g. *10m width*). | Area, elevation, and location | *2*<br>$^2$<br>^2<br>*across*<br>*above sea level*<br>*cell* | |
| | Length, diameter and width | *diam**<br>*wide*<br>*dbh*<br>*d.b.h.*<br>*dap*<br>*long*<br>*largo*<br>*comprimento*<br>*width*<br>*tall at anthesis* | When *in* or *de* was placed between a unit and an expression (e.g., *1m in width*), they were removed together. |
| | Others | *o.*<br>*o h*<br>*.o.*<br>*o*<br>*nv*<br>*v*<br>*sv* | Swedish or Norwegian representing values other than plant height. |

**Table S8**. Continued.

| Order of expression, number, and unit | Category of expression | Expression | Note |
|---|---|---|---|
| An expression is placed immediately **before** the combination of number and unit (e.g. *width 10m*). | Length, diameter, and width | *width*<br>*width to*<br>*width at*<br>*dbh of*<br>*dbh*<br>*d.b.h.*<br>*dap*<br>*diameter*<br>*diametro de*<br>*徑*<br>*長* | When = or *:* was placed between an expression and a number (e.g., *width = 1m*), they were removed together. |

**Table S9**. List of units used for plant height extraction. For eFloras, only *mm*, *cm*, *dm*, and *m* were considered.

| Unit | Resolved unit |
|---|---|
| *m* | meter |
| *meter* | |
| *metro* | |
| *metre* | |
| *met* | |
| *mt* | |
| *米* | |
| *公尺* | |
| *dm* | decimeter |
| *cm* | centimeter |
| *centimeter* | |
| *厘米* | |
| *公分* | |
| *mm* | millimeter |
| *foot* | foot |
| *feet* | |
| *ft* | |
| *'* | |
| *inch* | inch |
| *in* | |
| *''* (two single-quotes) | |
| *"* (one double-quote) | |
| *""* (two double-quotes) | |

**Text S1**: Patterns of expression used to extract of values of plant height (*PH*) from GBIF and eFloras. The definitions of the numbers, units, and their combinations are the same as those in Table S8. However, values with more than three digits (e.g., *1000*) were ignored.

For GBIF, we used two expression patterns to extract *PH* values. The first pattern, which detects expressions such as *10m tree* or *10 m tall* was

{Combination of numbers and units} {{*LS* or *GF*} or {*tall*/*high*/*alt*/*alt.*/*alltura*/*alto*/*altura*/*haut*/*in height*/*heigh*/*height*}},

where {*LS* or *GF*} represents *LS* or *GF*, as shown in Table S3 {*tall*/*high*/*alt*/*alt.*/*alltura*/*alto*/*altura*/*haut*/*in height*/*heigh*/*height*} are shown in parentheses.

The second pattern, which detected expressions such as *tree 10 m* or *height 10 m* was

{{*LS*, *GF*} or {whole plant body}} {*to*/*up to*/*de*/*ate*/*de ate*/*com*/*cerca de*/*de cerca de*/*com cerca de*/*of*/*:*/*,*} {Combination of number and unit},

where {*LS* or *GF*} is an expression representing *LS* or *GF* shown in Table S3, {whole plant body} is an expression representing the whole plant body shown in Table S10, and {*to*/*up to*/*de*/*ate*/*de ate*/*com*/*cerca de*/*de cerca de*/*com cerca de*/*of*/*:*/*,*} is one of the expressions in the brackets, which can also be missing (i.e., both *tree to 10 m* and *tree 10m* are possible).

For eFloras, the following pattern was used:

{*plant*/*culm*/*stem*/*herb*/*tree*/*shrub*/*vine*/*liana*/*climber*/*annual*/*biennial*/*perennial*} {Combination of numbers and units}

where the first bracket is an expression within the brackets.

**Table S10**. The list of expressions representing the whole plant body other than expressions of *LS* and *GF* in Table S3 was used for the extraction of *PH* from GBIF.

| Expression |
|---|
| *alltura* |
| *alt* |
| *alt.* |
| *alto* |
| *altura* |
| *climber* |
| *climbing* |
| *de h* |
| *erect* |
| *ereta* |
| *escandente* |
| *fern* |
| *haut* |
| *hauteur* |
| *height* |
| *(hight)* |
| *(high)* |
| *ht* |
| *ht.* |
| *hgt* |
| *hgt.* |
| *h* |
| *plant* |
| *planta* |
| *stem* |
| *straight* |
| *succulent* |
| *twiner* |
| *twining* |
| *株高* |
| *體高* |
| *高度* |
